# Neural tracking of the speech envelope predicts binaural unmasking

**DOI:** 10.1101/2024.05.22.595294

**Authors:** Benjamin Dieudonné, Lien Decruy, Jonas Vanthornhout

**Affiliations:** KU Leuven – University of Leuven, Department of Neurosciences, Experimental Oto-rhino-laryngology, Herestraat 49 bus 721, 3000 Leuven, Belgium

## Abstract

Binaural unmasking is the remarkable phenomenon that it is substantially easier to detect a signal in noise, when the interaural parameters of the signal are different from those of the noise – a mechanism that comes in handy in so-called cocktail party scenarios. In this study, we investigated the effect of binaural unmasking on neural tracking of the speech envelope. We measured EEG in 8 participants who listened to speech in noise at a fixed signal-to-noise ratio (−12 dB or −9 dB, depending on the speech material), in two conditions: one where speech and noise had the same interaural phase difference (both speech and noise having an opposite waveform across ears, *SπNπ*), and one where the interaural phase difference of the speech was different from that of the noise (only the speech having an opposite waveform across ears, *SπN*0). We measured a clear benefit of binaural unmasking in behavioral speech understanding scores, accompanied with increased neural tracking of the speech envelope. Moreover, analyzing the temporal response functions revealed that binaural unmasking also resulted in decreased peak latencies and increased peak amplitudes. Our results are consistent with previous research using auditory evoked potentials and steady-state responses to quantify binaural unmasking at cortical levels. Moreover, they confirm that neural tracking of speech is modulated by speech understanding, even if the acoustic signal-to-noise ratio is kept constant.

**Significance Statement:** Binaural unmasking in an important contributing factor to speech understanding in noisy environments. This is the first time that this benefit in speech understanding is measured objectively with EEG. From a clinical perspective, these results could enable the evaluation of binaural speech understanding mechanisms in populations for whom behavioral measures are difficult to obtain, such as young children or people with cognitive impairment. Moreover, behavioral research has shown that many pathologies (e.g., asymmetric hearing loss, auditory neuropathy, and age-related deficits) are more sensitive to tests that contain a binaural processing component. Our paradigm could enable the detection of such pathologies with an objective approach using neural tracking of speech.

## 1 Introduction

People are remarkably good at recognizing speech when multiple speakers are talking simultaneously, in so-called “cocktail party” scenarios (Cherry, 1953; Bronkhorst, 2000). One important contributing factor is that hearing with two ears – binaural hearing – provides a substantial benefit in detecting a target signal in noise (Bronkhorst and Plomp, 1988; Dieudonnéand Francart, 2019). In particular, when the interaural parameters of the signal differ from those of the noise, it becomes substantially easier to detect or identify the signal. For example, when identical tones and noise are presented to both ears, detectability is improved by up to 15 dB simply by inverting the phase of the tone or the noise in one ear (Hirsh, 1948). This benefit is referred to as the binaural masking level difference (BMLD), and the effect is called binaural unmasking.

Binaural unmasking is presumed to occur as a neural subtractive mechanism, and strongly relies on the integrity of phase information transmitted by the auditory brainstem (Moore, 1991; Palmer and Shackleton, 2002; Ishida and Stapells, 2009; Culling and Lavandier, 2021). The increase in signal detectability due to interaural differences has been measured at the neuronal level in the inferior colliculus (midbrain) and auditory cortex in studies with guinea pigs (Caird et al., 1991; McAlpine et al., 1996; Jiang et al., 1997; Gilbert et al., 2015). In humans, binaural unmasking has also been measured in many (non-invasive) electrophysiological studies using electroencephalography (EEG) (Edwards et al., 1971; Fowler, 2017). Interestingly, binaural unmasking effects are most consistent in studies that measured responses at the cortical level (e.g., an increase in peak amplitudes and decrease in peak latencies of the auditory evoked potential (AEP)) (Kevanishvili and Lagidze, 1987). At subcortical levels (auditory midbrain and auditory brainstem), results differ across studies. For example, Kevanishvili and Lagidze (1987) were only able to measure unmasking effects at a cortical level, while they found no effects at the auditory brainstem (wave V response) nor auditory midbrain (mid-latency response peak Na). On the other hand, Wilson and Krishnan (2005) and Clinard et al. (2017) found an effect of binaural unmasking on frequency-following responses, which they presumed to originate from the auditory brainstem.

While binaural unmasking of speech has been measured behaviorally already in the 1940s (Licklider, 1948), electrophysiological studies have only shown binaural unmasking for rudimentary stimuli, such as click trains or (modulated) tones in noise (for an overview, see Fowler, 2017). These stimuli allow averaging trials or frequency-domain analysis to overcome the noisy character of EEG. However, more recent analysis methods allow measuring electrophysiological responses to single-trial, continuous speech as well (Lalor et al., 2009; Ding and Simon, 2012; Brodbeck and Simon, 2020). The approach involves decoding speech features from EEG recordings (backward modeling) or predicting EEG responses to specific speech features (forward modeling). For example, the speech envelope (an important acoustic cue for speech understanding, see Shannon et al., 1995) can be reconstructed from EEG measured during speech listening. The reconstruction accuracy provides insights into the cortical tracking of speech, and can even predict speech understanding (Vanthornhout et al., 2018; Iotzov and Parra, 2019; Etard and Reichen-bach, 2019; Lesenfants et al., 2019; Gillis et al., 2022). Moreover, when predicting the EEG response from the speech envelope (forward modeling), the (linear) prediction model – called temporal response function (TRF) – reflects how the EEG response is modulated by the envelope at different time shifts. As such, it can be interpreted as an AEP from which neural response amplitudes and latencies can be inferred.

In this study, we aim to measure the effect of binaural unmasking on neural tracking of continuous speech. We hypothesize that binaural unmasking will increase envelop reconstruction accuracies, as we expect that the improved speech understanding goes hand in hand with an improved envelope representation at the cortical level. We also hypothesize that binaural unmasking will increase TRF peak amplitudes, and decrease TRF peak latencies, in accordance with studies on AEPs (Kevanishvili and Lagidze, 1987).

## 2 Materials and methods

### 2.1 Participants

We tested 8 young female normal-hearing listeners, that is, aged between 18 and 27 years old and having pure-tone hearing thresholds better than 20 dB HL at all octave audiometric frequencies from 250 to 8000 Hz. This study was approved by the Medical Ethics Committee Research UZ/KU Leuven (project number S57102). All participants gave written informed consent for their participation in the study.

### 2.2 Experimental design

#### 2.2.1 Experimental set-up

The experiment took place in a triple-walled, soundproof booth, that was equipped with a Faraday cage to avoid electromagnetic interference. Stimuli were presented through ER-3A insert phones (Etymotic Research, Elk Grove Village, IL) via an RME Hammerfall DSP Multiface soundcard (Audio AG, Haimhausen, Germany), using the software platform APEX 3 (Francart et al., 2008). We recorded EEG using a BioSemi ActiveTwo system (Amsterdam, The Netherlands), with 64+2 active electrodes, at a sampling frequency of 8192 Hz.

#### 2.2.2 Stimuli

We used two stimulus types as target speech: a Matrix sentence test (Luts et al., 2015) and continuous narrated stories. All were spoken in Flemish (Dutch), the native language of our participants.

The Matrix sentence test consists of 13 lists of 20 sentences uttered by a female speaker. Each sentence lasts about 2 seconds and has the same grammatical structure (name, verb, numeral, adjective, object). The Matrix test allows for a reliable behavioral measurement of binaural unmasking.

We used three stories, all uttered by a male speaker: one story (*Story0* around 14 minutes) to verify neural tracking in silence, and two stories to measure binaural unmasking (*Story1* around 16 minutes, and *Story2* around 11 minutes). We used stories to verify whether binaural unmasking could also be measured in a more realistic listening condition.

As masking noise, we used stationary speech-weighted noise with the same long-term average speech spectrum as the stimuli.

Speech was always presented at 55 dBA, while the signal-to-noise ratio (SNR) was set via the noise level. As such, the speech presentation was always exactly the same across conditions.

#### 2.2.3 Conditions

We presented speech in noise in two conditions, as represented in Figure 1: *SπNπ* and *SπN*0.

**Figure 1.**
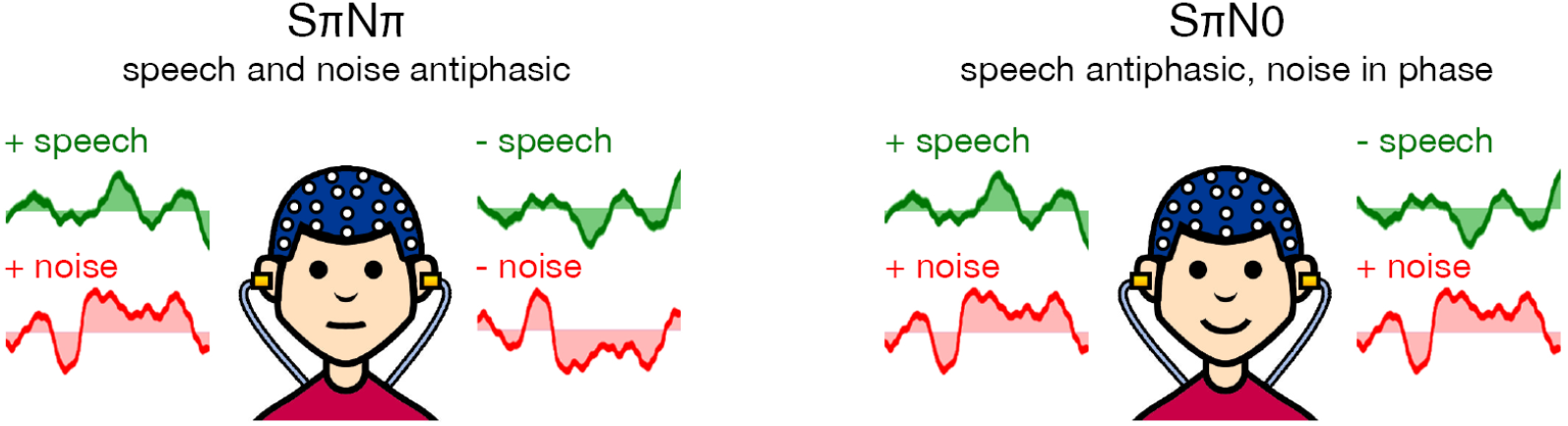
Graphical representation of the two listening conditions. For the left condition (*SπNπ*), both speech and noise have the same IPD of 180°. For the right condition (*SπN*0), the speech has an IPD of 180°, and the noise has no IPD. When presenting the same signal-to-noise ratio in both conditions, higher speech intelligibility is expected in the right condition (*SπN*0) due to binaural unmasking.

- For *SπNπ*, both speech and noise have an interaural phase difference (IPD) of 180° (that is, the signal presented to the right ear is exactly the same as in the left ear, except for a minus sign). When speech and noise have the same IPD, no binaural unmasking is expected, making *SπNπ* the difficult listening condition.
- For *SπN*0, only the speech has an IPD of 180°, and the noise has no IPD. When speech and noise have a different IPD, binaural unmasking is expected, making *SπN*0 the easier listening condition.

Note that there are many potential condition-pairs to measure binaural unmasking, but comparing *SπNπ* with *SπN*0 should result in the largest binaural unmasking effects (Robinson and Jeffress, 1963), while also keeping the interaural phase difference of the speech constant across conditions.

#### 2.2.4 Procedure

The experiment started with a brief behavioral training phase to get used to the Matrix speech material. In this training phase, we measured the speech reception threshold (SRT) (the SNR at which 50% of speech could be understood) with an adaptive procedure (Brand and Kollmeier, 2002). We did this for 3 lists of the condition *SπNπ*, and 1 list for the condition *SπN*0.

The EEG experiment consisted of 4 parts, that were randomized in order for each participant:

##### 1 Matrix sentences

we measured percentage of correctly understood words behaviorally for 2 lists for condition *SπNπ*, and 2 lists for the condition *SπN*0, at a fixed SNR of −12 dB^1^. Each list consisted of 20 sentences with 5 words per sentence. The order of lists was randomized for each participant.

##### 2 Story0

we presented this story without noise and diotically (that is, without IPD) to verify whether we could measure neural tracking for each participant. We were able to measure significant neural tracking in each of our participants for this condition; these results are left out.

##### 3 Story1

the story was cut in 4 fragments of approximately equal length, with a condition assigned to each fragment (2 times *SπNπ*, 2 times *SπN*0, in randomized order). Before presenting the story, the participant got a short introduction to the context of the story. After every fragment, we asked the participant for a subjective rating of speech understanding (“How much of the story have you understood (in percent)?”), and a question about content of story. The story was presented at a fixed SNR of −9 dB (from pilot tests, we expected the unmasking effect to be the strongest around this SNR for the story, avoiding ceiling or floor effects).

##### Story2

same as **Story1**.

### 2.3 EEG analyses

#### 2.3.1 Pre-processing

Before the actual EEG analyses, we performed the following pre-processing steps:

1. remove drift with a zero-phase highpass Butterworth filter of order 1 with cut-off at 0.1 Hz
2. downsample from 8192 Hz to 128 Hz to reduce processing time (using Matlab’s “resample” function which applies zero-phase anti-aliasing)
3. remove eyeblink artefacts via a multi-channel Wiener filter (Somers et al., 2018)
4. re-reference EEG to the common average
5. filter the EEG between 1 Hz and 8 Hz (this frequency band has shown to be relevant for speech understanding (Ding and Simon, 2014)) with a second-order Butterworth filter (in the forward and reverse direction to have zero phase distortion, which means that the actual filter order is 4)
6. normalise the EEG channels to unit variance and zero mean

#### 2.3.2 Calculation of the actual acoustic speech envelope

We chose the acoustic speech envelope as the speech feature in our EEG analysis. The speech envelope *s*(*t*) was calculated by filtering the speech stimulus (without noise) through a gammatone filterbank with 28 filters spaced by equivalent rectangular band-width with center frequencies of 50 Hz to 5 kHz (Søndergaard and Majdak, 2013), calculating the envelope in each band by taking the absolute value, compressing each envelope by raising it to the power of 0.6, then averaging the resulting narrow-band envelopes to get a broadband envelope (Biesmans et al., 2016). Then we performed similar pre-processing steps as for the EEG:

1. apply same filter that was used to remove drift from EEG
2. downsample from 48 000 Hz to 128 Hz
3. filter between 1 Hz and 8 Hz with a second-order Butterworth filter (in the forward and reverse direction to have zero phase distortion, which means that the actual filter order is 4)
4. normalise the envelope to unit variance and zero mean

Note that inverting the phase of the speech stimulus (see section 2.2.3) has no effect on its envelope.

#### 2.3.3 Backward model (stimulus envelope reconstruction)

We measured neural tracking of the speech envelope by calculating the Spearman correlation between the actual acoustic speech envelope *s*(*t*), and the envelope *ŝ*(*t*) that we reconstructed from the EEG that was measured during stimulus presentation (Vanthornhout et al., 2018).

The reconstructed stimulus envelope *ŝ*(*t*) was calculated by applying a linear decoder *g*(*n, τ*) to the (pre-processed) EEG response *r*(*t, n*), with *t* referring to time, *n* referring to the EEG channel and *τ* the latency of the brain response (Crosse et al., 2016):

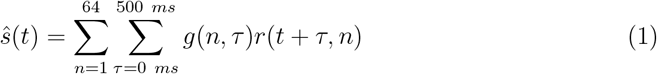

or in matrix notation:

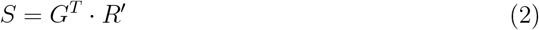

with *R*^t^ the matrix of the time-shifted EEG with dimensions (channels * latencies, time), *S* the matrix of the envelope with dimensions (1, time), and *G* the decoder matrix with dimensions (channels * latencies, 1).

Thus, the linear decoder is a spatio-temporal filter that combines the EEG channels (1 to 64) and their time-shifted versions (post-stimulus integration window from 0 to 500 ms) to optimally reconstruct the envelope. The coefficients can be found using ridge regression on a training data set:

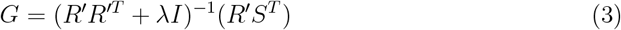

with λ the ridge parameter (regularization parameter) set to the maximum absolute value of the auto-correlation matrix *R*^′^*R*^′*T*^ of the time-shifted EEG, and *I* the identity matrix. For each participant, for each listening condition (Matrix, Story1 and Story2), a (subject-dependent and condition-dependent) linear decoder was calculated and applied in a cross-validated way:

- For the matrix sentences, we combined the data of the two lists corresponding to each condition (*SπN*0 and *SπNπ*). For training the decoder, we set the parts of the envelope that contained participant’s answer to zero. We split the data in 40 folds (1 fold per sentence), then reconstructed the envelope from the EEG per fold based on a decoder trained on all other folds. These 40 reconstructed speech envelopes (1 per sentence) are then concatenated to obtain a reconstructed envelope *ŝ*(*t*) per condition. The silences of the actual envelope and reconstructed envelope are removed before calculating the Spearman correlation.
- For the stories *Story1* and *Story2*, we combined the data of the two fragments corresponding to each condition (*SπN*0 and *SπNπ*). We split the data in 40-second folds (about 7 to 12 folds per story per condition), then reconstructed the envelope from the EEG per fold based on a decoder trained on all other folds. The reconstructed speech envelopes (1 per fold) are again concatenated to obtain a reconstructed envelope *ŝ*(*t*) per condition per story.

#### 2.3.4 Forward model (TRF analysis)

##### Calculation of the TRF

The TRF reflects how the EEG response *R*(*t, n*) (electrode *n*) is modulated by the envelope *s*(*t*) at different time-shifts *τ* (pre-response integration window from 0 to 500 ms). Mathematically, the model predicts a reconstructed EEG response 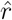 (*t, n*) (Crosse et al., 2016):

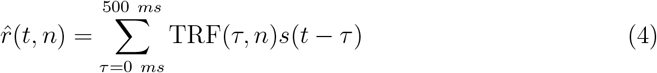

or in matrix notation (for each electrode *n* separately):

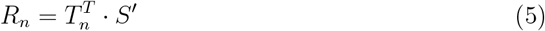

with *S*^′^ the matrix of the time-shifted envelope with dimensions (latencies, time), *R*_*n*_ the matrix of the EEG at channel *n* with dimensions (1, time), and *T*_*n*_ the TRF matrix with dimensions (latencies, 1).

The TRF coefficients can be found using regularized regression, as described by Crosse et al. (2016):

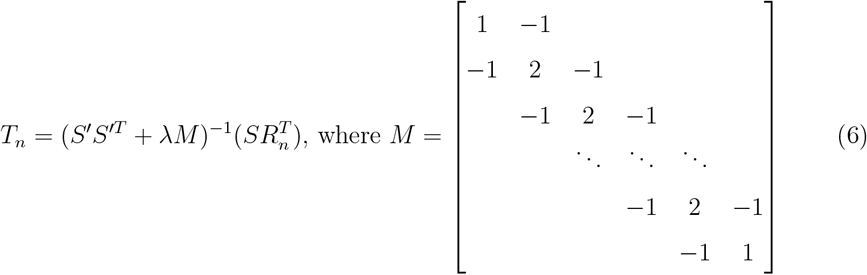

with λ the regularization parameter set to the maximum absolute value of the auto-correlation matrix *S*^′^*S*^′*T*^ of the time-shifted stimulus envelope.

For the forward model, we combined the data of the two stories (per condition, per participant), and did no analysis on the data from the matrix sentences. As such, we could calculate a TRF for each condition, for each participant, for each EEG electrode. As we were only interested in the TRF itself, and not in the EEG reconstruction accuracies, each TRF was calculated once using all corresponding data (i.e., without cross-validation).

##### Averaging the TRFs over electrodes

After calculating the TRFs for each electrode (per participant, per condition), we matched the polarity of the TRFs over electrodes in the following way:

1. Calculate the EEG reconstruction accuracy (in a cross-validated way, similar to the backward model) for each electrode, for each subject, for each condition. From this, we found that P9 had the highest average reconstruction accuracy.
2. For each electrode, calculate the average TRF (over subjects and conditions). Then, calculate the correlation of this average TRF with the TRF from the electrode that has the highest average reconstruction accuracy (electrode P9). If this correlation is negative, the sign of the TRFs of that electrode (for each subject, for each condition) is switched.

Then, we calculated a single TRF per subject per condition by averaging over all (polarity-matched) electrodes. Finally, the sign of the resulting average TRFs is switched one more time, such that the peaks have the expected polarity (P1, P2 positive; N1, N2 negative). The resulting average TRF (for each subject, for each condition) is equivalent to the result of applying spatial filter with only +1s and −1s as filter coefficients, which is depicted in the topoplot in Figure 3.

##### TRF peak detection

Finally, we calculated the peak amplitudes and latencies of the average TRFs for each subject, for each condition, and also for the TRFs averaged over subjects, per condition. The peaks were simply detected by finding the first two local maxima (P1 and P2) and first two local minima (N1 and N2) in the TRF (only considering latencies greater tha*NO*). To increase temporal resolution for the peak latencies (about 8 ms at a sampling rate of 128 Hz), we fitted a parabola through each extremum and its two neighbouring values, and used the amplitude and latency of the extremum of the resulting parabola.

### 2.4 Statistical analysis

All signal processing described above was performed using MATLAB (version R2023b).

Statistical analyses were done with R (programming language). For each of the following outcomes, we tested the difference across subjects using an exact one-sided Wilcoxon signed-rank test (performed using the coin package by Hothorn et al., 2008):

- behavioral speech understanding score (separately for *Story*1, *Story*2, and matrix sentences)
- neural envelope tracking (backward model reconstruction accuracy; separately for *Story*1, *Story*2, and matrix sentences)
- TRF peak amplitude (P1, N1, P2, N2; *Story*1 and *Story*2 combined)
- TRF peak latency (P1, N1, P2, N2; *Story*1 and *Story*2 combined) The significance level was set at *α* = 0.05.

## 3 Results

### 3.1 Backward model (stimulus envelope reconstruction)

The results are shown in Figure 2 (behavioral scores in the top row, neural tracking in the bottom row).

**Figure 2.**
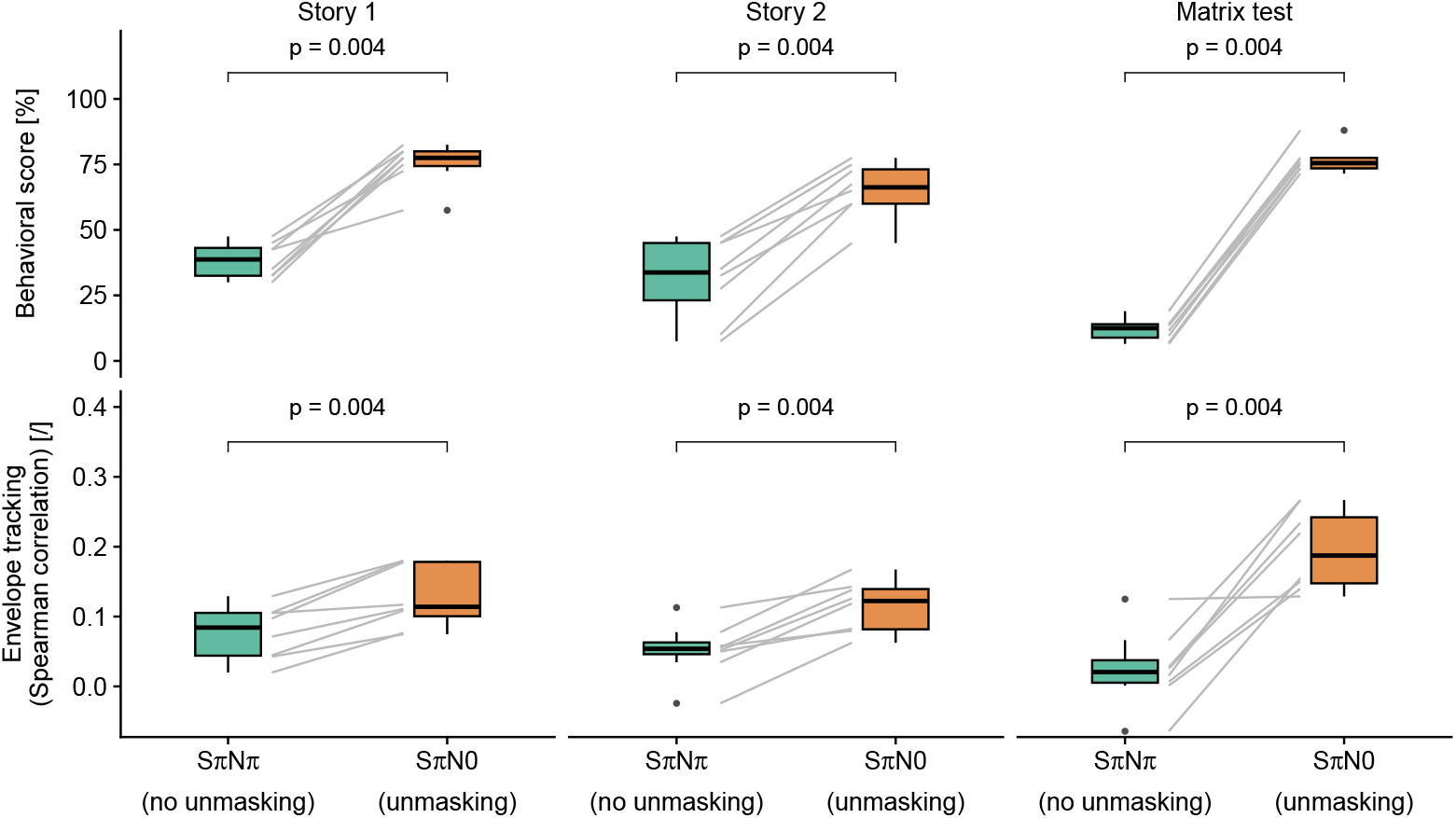
Comparison of behavioral scores (top row) and backward model stimulus reconstruction accuracies (bottom row). We find consistent significant effects of binaural unmasking: for all subjects and listening conditions, we found better speech understanding scores in the *SπN*0 condition than in the *SπNπ* condition, and higher neural tracking scores in the *SπN*0 condition than in the *SπNπ* condition.

**Figure 3.**
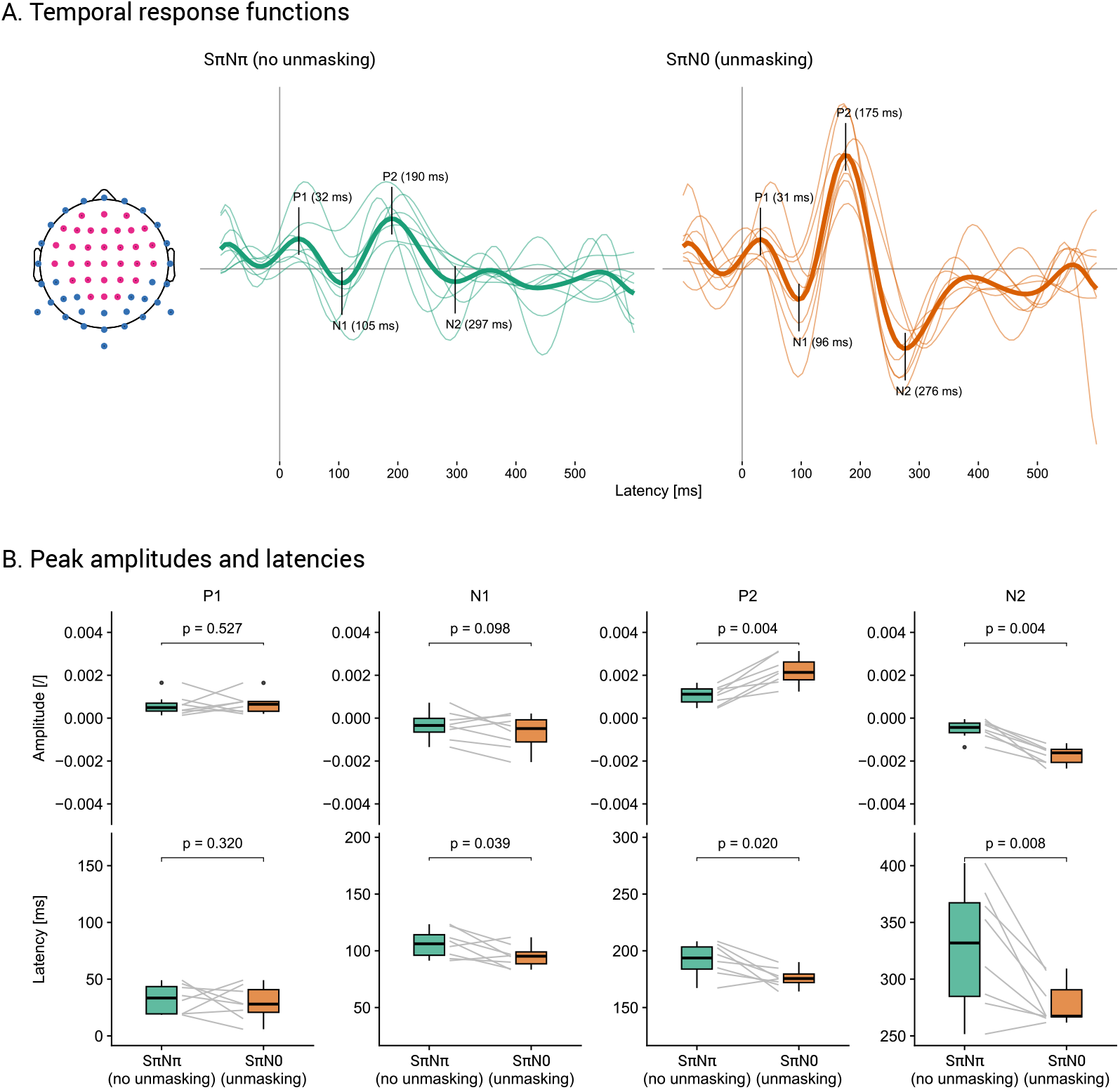
Results of the forward modelling analyses, using the combined EEG recordings from Story1 and Story2. For N1, P2 and N2, we find a significant decrease in peak latency due to binaural unmasking; for P2 and N2, we also find a significant increase in peak amplitude due to binaural unmasking.

As expected, we measured higher behavioral speech understanding scores in the *SπN*0 condition than in the *SπNπ* condition due to binaural unmasking (*Z* = 2.52, *p* = 0.004 for all pairwise comparisons). The effect was consistent for all participants (each colored line on the figure represents the scores of 1 participant), and for each speech material – even for the subjective speech understanding estimates for the stories.

The neural tracking results agreed well with the behavioral results: we measured higher neural tracking scores in the *SπN*0 condition than in the *SπNπ* condition due to binaural unmasking (*Z* = 2.52, *p* = 0.004 for all pairwise comparisons). The effect was again consistent for all participants, and for each speech material.

### 3.2 Forward model (TRF analysis)

The results are shown in Figure 3 (spatial filter and TRFs in panel A, TRF peak characteristics in panel B).

As visualized in the topoplot (Figure 3A, left side), the TRFs (Figure 3A, right side) are an average of the TRFs over all electrodes, with the central electrodes having the original sign, and the peripheral electrodes having a switched sign – to ensure that all TRFs have the same polarity before averaging.

By visual inspection of the TRFs averaged over subjects per condition (bold lines on Figure 3A, right side), it can be seen that we measured larger (amplitude) and earlier (latency) TRF peaks in the *SπN*0 condition than in the *SπNπ* condition due to binaural unmasking, with larger differences for the later peaks (P2 and N2).

When investigating the individual peaks over subjects (Figure 3B), we found significant differences in the latencies for N1 (*Z* = −1.82, *p* = 0.039), P2 (*Z* = −2.10, *p* = 0.020), and N2 (*Z* = −2.38, *p* = 0.008), and significant differences in the amplitudes for P2 (*Z* = 2.52, *p* = 0.004) and N2 (*Z* = −2.52, *p* = 0.004). We found no significant difference in peak latency for P1 (*Z* = −0.56, *p* = 0.320), nor in peak amplitude for P1 (*Z* = 0.00, *p* = 0.527) and N1 (*Z* = −1.400, *p* = 0.098).

A. Individual and group-averaged TRFs for the two listening conditions (averaged across electrodes, where we switched the sign of the TRFs from the blue electrodes on the topoplot before averaging). Note that the two figures share the same arbitrary unit for the amplitude on the vertical axis.
B. Peak amplitudes (top row) and latencies (bottom row) for extracted P1, N1, P2 and N2 peaks from the individual TRFs.

## 4 Discussion

In this study, we measured the effect of binaural unmasking on neural tracking of the speech envelope. Using a backward model, we found a consistent increase in envelope reconstruction accuracies due to binaural unmasking. Using a forward model, we found an increase in TRF peak amplitudes (P2 and N2), and decrease in TRF peak latencies (N1, P2 and N2) due to binaural unmasking. However, we found no significant effect on TRF peak amplitude and peak latency of P1, nor a significant effect on the TRF peak amplitude of N1.

Except for the lack of significant effects on the earliest cortical peaks, the neural tracking results were in line with our hypotheses, and agreed well with the behavioral results. It’s worth noting that the effect of binaural unmasking could also be evaluated subjectively, by asking participants about their comprehension of the stories for the different conditions. The outcome was consistent across all participants and both stories, aligning with previous research findings that subjective assessments of understanding of continuous natural speech can provide reliable results (Decruy et al., 2018).

### 4.1 Electrophysiological measurements of binaural hearing

The binaural system is responsible for two main tasks: localization of sound sources in the horizontal plane and improved speech understanding in noise. The former had already been measured in neural tracking paradigms, by relating sound source trajectories to EEG (Bednar and Lalor, 2018, 2020). We have now shown that neural tracking of speech can also predict binaural benefits of speech understanding in noise.

As mentioned in the introduction, the improved detectability of sounds due to binaural hearing had already been measured in EEG studies long time ago (Kevanishvili and Lagidze, 1987). To our knowledge, this is the first time that binaural unmasking is measured via neural tracking of continuous speech. We found results that are in accordance with cortical response studies: (1) we found increased backward model correlations with binaural unmasking, in accordance with higher low-frequency auditory steady state response (ASSR) response amplitudes (Wong and Stapells, 2004), and (2) we found increased peak amplitudes and decreased peak latencies with binaural unmasking, in accordance with studies on AEPs (Kevanishvili and Lagidze, 1987).

However, in contrast with the study of Kevanishvili and Lagidze (1987), we found no significant effect on the peak latency of P1, and no significant effect on the peak amplitudes of P1 and N1. Although we did not report it above, we also did not find a significant effect when considering the P1N1 difference (as in the study by Kevanishvili and Lagidze, 1987) instead of the individual P1 and N1 amplitudes. The lack of effect on N1 amplitude could be driven by a lack of statistical power (low sample size). From the individual TRFs (Figure 3A), it can also be seen that the earlier peaks (P1 and N1) are less well defined than the later peaks (P2 and N2), which might be resolved with longer within-subject EEG recordings. Lastly, given the latency of the P1 peaks (around 30 ms), these P1 peaks might actually be Pa peaks or a mixture of Pa and P1. A lack of effect of binaural unmasking on Pa peaks would be in accordance with the AEP study by Kevanishvili and Lagidze (1987).

Kevanishvili and Lagidze (1987) concluded from their results (i.e., binaural unmasking can only be measured objectively in late cortical responses) that the masking level difference is operated at the cortical level. Although our results could lead to the same conclusion, one has to keep in mind that binaural unmasking has also been measured at the neuronal level in the inferior colliculus (auditory midbrain) in animal models (Caird et al., 1991; McAlpine et al., 1996; Jiang et al., 1997), and on frequency-following responses (presumed to originate from the auditory brainstem) in humans (Wilson and Krishnan, 2005; Clinard et al., 2017).

In future research, it would be interesting to verify whether the effect of binaural unmasking on earlier neural responses could be replicated in a study with continuous speech. Indeed, novel signal processing methodologies allow to measure early subcortical responses (5 to 15 ms) to continuous speech (Forte et al., 2017; Van Canneyt et al., 2021b,a; Kulasingham et al., 2024). These studies employ similar backward and forward modelling approaches as applied in this study, but use the fundamental frequency of the voice instead of slow envelope modulations as a speech feature. However, even in quiet listening conditions, these subcortical responses are typically very small compared to the cortical responses that we discussed in this paper. Therefore, in a preliminary analysis we found that our dataset is not suitable for such an early-response analysis. To our knowledge, such early subcortical responses to continuous speech have not yet been measured in challenging (i.e,. noisy) listening conditions.

### 4.2 Consequences for neural envelope tracking research

There is an ongoing debate on what neural tracking of acoustic speech features actually measures. Quite some research has shown that there is a clear link between neural envelope tracking and speech understanding (Ding and Simon, 2013; Etard and Reichenbach, 2019; Vanthornhout et al., 2018; Riecke et al., 2018; Iotzov and Parra, 2019). However, these studies typically modulate speech understanding by adapting the SNR. When speech understanding is modulated in different ways – e.g., by changing the speech rate, or by degrading the spectral resolution with vocoders – neural tracking of acoustic features is not always the best predictor of speech understanding (Verschueren et al., 2022; Karunathilake et al., 2023; Kösem et al., 2023). In this study, we modulated speech understanding yet in another way: we kept the acoustic SNR constant, and merely switched the phase of the noise in one of the ears. In the perspective of neural tracking research, it is remarkable that we observed such a large increase in reconstruction accuracies without having to increase the acoustic SNR. Neural tracking of speech appears to measure at least the representation (or even SNR) of the acoustic envelope in the auditory cortex after the binaural system has denoised the signal, and therefore agrees well with speech intelligibility in this experimental paradigm.

Considering the forward model, the effect of improved intelligibility (due to binaural unmasking) agrees well with the literature as well: Muncke et al. (2022) and Chen et al. (2023) also measured increased TRF peak amplitudes and decreased TRF latencies with improved intelligibility.

Finally, envelope reconstruction has also been suggested as a way to measure auditory attention. When a person listens to multiple sound sources, the envelope of the attended sound source will have a better reconstruction accuracy than the unattended streams (O’sullivan et al., 2015). In a two-speaker paradigm, it has been found that using realistic spatial cues (interaural time and level differences) results in larger reconstruction accuracies than when each speaker is presented to a separate ear without spatial cues (Das et al., 2016). This could potentially be explained as a better cortical representation of the speakers due to binaural unmasking.

### 4.3 Clinical relevance

Neural tracking of speech has the potential to measure speech understanding in populations where conventional methods are impractical, such as young children, individuals with dementia, coma, aphasia, etc. (Gillis et al., 2022; Van Hirtum et al., 2023b,a). The fact that it can also robustly measure binaural processing (using the backward model, we measured a consistent binaural unmasking effect on a single-subject level across speech materials!) opens a whole new window of diagnostic opportunities.

Many pathologies are indeed more sensitive to tests that contain a binaural processing component (Olsen et al., 1976). De Sousa et al. (2020) have shown that (symmetric and asymmetric) hearing losses can be detected with higher sensitivity with a speech understanding test using antiphasic noise. Auditory neuropathy also has a stronger (negative) effect on speech understanding scores in conditions where binaural processing might have helped (Rance et al., 2012). Slight hearing losses, age-related deficits, and even suprathreshold deficits have shown to affect binaural unmasking effects (Bernstein and Trahiotis, 2016; Tolnai et al., 2024; Eddins and Eddins, 2018; Anderson et al., 2018). It is expected that these pathologies affect the binaural unmasking benefit in neural tracking of the speech envelope – this remains to be investigated in future research.

## 5 Acknowledgments

We would like to thank professor Tom Francart for his insightful comments on the experimental protocol, the data analyses and the manuscript. We also thank Jolien Smeulders, Sara Peeters, and Merel Dillen for their help with the data collection, and our participants for their patience and enthusiasm during our experiment.

This research is funded by grants of the Research Foundation Flanders (FWO) for Jonas Vanthornhout (PhD grant JV: 1S10416N postdoc grant JV: 1290821), and Benjamin Dieudonné(Phd grant BD: 1S45817N).

## Footnotes

1 To be able to measure binaural unmasking in the constant procedure at an SNR of −12 dB, the SRT for *SπNπ* (difficult) should be higher (worse) than −12 dB, while the SRT for *SπN*0 (easier) should be lower (better) than −12 dB. This was the case for all our participants, as verified with the results from the behavioral training phase.

